# Structure of nucleosome-bound human PBAF complex

**DOI:** 10.1101/2022.05.20.492795

**Authors:** Li Wang, Jiali Yu, Zishuo Yu, Qianmin Wang, Wanjun Li, Yulei Ren, Zhenguo Chen, Shuang He, Yanhui Xu

**Affiliations:** Fudan University Shanghai Cancer Center, Institutes of Biomedical Sciences, State Key Laboratory of Genetic Engineering and Shanghai Key Laboratory of Medical Epigenetics, Shanghai Medical College of Fudan University, Shanghai 200032, China; The International Co-laboratory of Medical Epigenetics and Metabolism, Ministry of Science and Technology, China, Department of Systems Biology for Medicine, School of Basic Medical Sciences, Shanghai Medical College of Fudan University, Shanghai 200032, China; Human Phenome Institute, Collaborative Innovation Center of Genetics and Development, School of Life Sciences, Fudan University, Shanghai 200433, China; The Fifth People’s Hospital of Shanghai, Shanghai Institute of Infectious Disease and Biosecurity, Shanghai Key Laboratory of Medical Epigenetics, and Institutes of Biomedical Sciences, Fudan University, Shanghai 200032, China

## Abstract

BAF and PBAF are mammalian SWI/SNF family chromatin remodeling complexes that possess multiple histone/DNA-binding subunits and create nucleosome-depleted/free regions for transcription activation. Despite structural studies of nucleosome-bound human BAF and yeast SWI/SNF family complexes, it remains elusive how PBAF-nucleosome complex is organized. Here we determined structure of 13-subunit human PBAF in complex with acetylated nucleosome in ADP-BeF_3_-bound state. Four PBAF-specific subunits work together with nine BAF/PBAF-shared subunits to generate PBAF-specific modular organization, distinct from that of BAF at various regions. PBAF-nucleosome structure reveals six histone-binding domains and four DNA-binding domains/modules, the majority of which directly bind histone/DNA. This multivalent nucleosome-binding pattern, not observed in previous studies, suggests that PBAF may integrate comprehensive chromatin information to target genomic loci for function. Our study reveals molecular organization of subunits and histone/DNA-binding domains/modules in PBAF-nucleosome complex and provides a framework to understand chromatin targeting of SWI/SNF family complexes.

## Introduction

The adenosine triphosphate (ATP)-dependent chromatin remodeling complexes (also known as remodelers) regulate chromatin architecture by reorganizing nucleosome positioning and content (Clapier and Cairns, 2009; Clapier et al., 2017). SWI/SNF is the prototype chromatin remodeler possessing nucleosome sliding activity and unique histone ejection activity, and by which creates nucleosome-depleted or nucleosome-free regions (NDRs/NFRs) on gene promoters required for transcription initiation (Boeger et al., 2004; Cairns et al., 1994; Cote et al., 1994). SWI/SNF complexes exist in SWI/SNF and RSC complexes in yeast and their counterparts in mammals are BRG1/BRM-associated factor (BAF) and polybromo-associated BAF (PBAF) complexes (Mashtalir et al., 2018; Wang et al., 1996; Xue et al., 2000), respectively. In line with their functional importance in gene regulation, BAF and PBAF are among the most frequently mutated complexes in cancer, as up to 20% of malignancies have alterations on coding genes of BAF/PBAF subunits (Kadoch and Crabtree, 2015; Kadoch et al., 2013; Mittal and Roberts, 2020; Wilson and Roberts, 2011).

BAF and PBAF are megadalton multi-subunit complexes, which share a catalytic subunit SMARCA4 (BRG1) and eight common auxiliary subunits, including ACTB, ACTL6A, BCL7A, SMARCB1 (BAF47), SMARCD1 (BAF60A), SMARCE1 (BAF57), and two SMARCC1/2 (BAF155 and BAF170, equivalent and termed SMARCC for simplicity) (Fig. 1a). The two complexes are distinguished by BAF-specific subunits ARID1A/B (BAF250A/B), DPF1/2/3 (BAF45B/D/C), and SS18 and PBAF-specific subunits ARID2 (BAF200), PHF10 (BAF45A), PBRM1 (BAF180), and BRD7. Despite recent advances in structural studies of nucleosome-bound human BAF (He et al., 2020; Mashtalir et al., 2020), yeast RSC (Patel et al., 2019; Wagner et al., 2020; Ye et al., 2019), and yeast SWI/SNF (Han et al., 2020) complexes, structure of human PBAF remains unknown. Consistent with the difference in complex composition, BAF and PBAF exhibit distinct preference in genomic localization (Alver et al., 2017; Michel et al., 2018) and functions (Kadoch and Crabtree, 2015; Mashtalir et al., 2021; Mittal and Roberts, 2020; Wilson and Roberts, 2011), suggesting a PBAF-nucleosome structure distinct from that of BAF-nucleosome. SWI/SNF family complexes consist of multiple (over 20 in PBAF) histone-binding and DNA-binding domains that are believed to facilitate genomic targeting and/or regulate remodeling activities of these complexes. However, it remains largely unknown how these histone/DNA-binding domains are organized within apo or nucleosome-bound SWI/SNF complexes.

**Figure 1.**
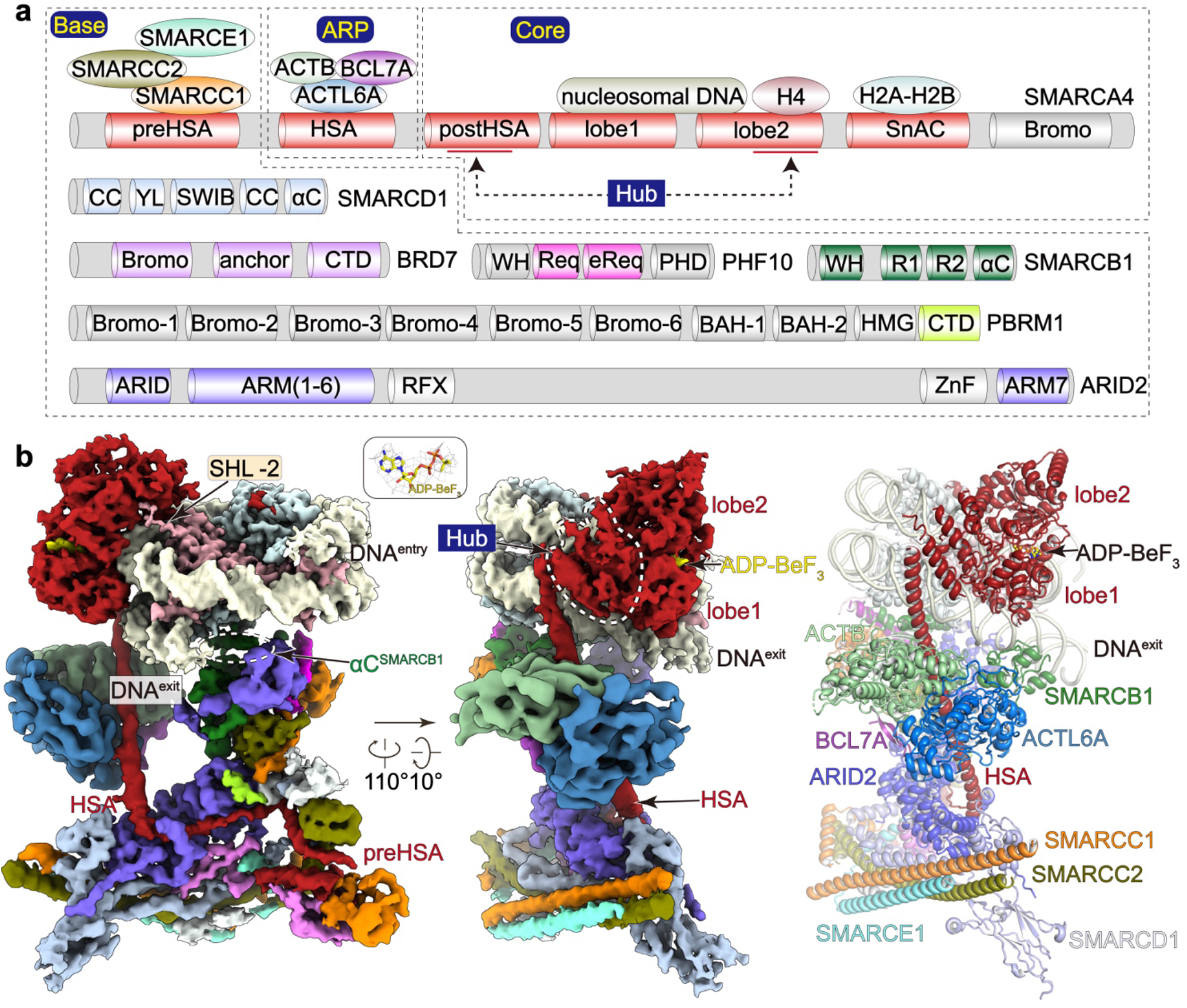
Cryo-EM structure of the nucleosome-bound PBAF. **(a)** Schematic diagram of the 13-subunit PBAF complex organized into the Base, ARP and Core modules. Histone/DNA-binding domains of indicated subunits are shown. Color scheme is used throughout figures if not elsewhere specified. **(b)** Composite cryo-EM map and structural model of PBAF-NCP in two different views. Close-up views of ADP-BeF_3_ is shown in cryo-EM map (mesh) and structural models (stick).

## Results

### Overall structure of the nucleosome-bound PBAF complex

We overexpressed the 13-subunit human PBAF complex in Expi293F suspension cells through co-transfection of plasmids containing full-length open reading frames (ORFs) of the catalytic subunit SMARCA4 and twelve auxiliary subunits (Fig. 1a, Extended Data Fig. 1a). The complex was purified to homogeneity for biochemical and structural analyses. In vitro chromatin remodeling assay showed that the purified PBAF converted a center-positioned nucleosome core particle (NCP) to three products, the end-positioned nucleosome, end-positioned tetrasome, and free DNA, indictive of its activity in sliding and ejection of nucleosome in a time-dependent manner (Extended Data Fig. 1b).

The chromatin substrates in cells of BAF/PBAF commonly contain acetylation at multiple sites of histone tails, which may regulate chromatin remodeling activity of SWI/SNF family complexes and facilitate their binding to nucleosome (Agalioti et al., 2002; Chatterjee et al., 2011; Mashtalir et al., 2021). We reconstituted unmodified nucleosome and performed an in vitro acetylation reaction using a mixture of two predominate human histone acetyltransferases (HATs), p300 and SAGA acetyltransferase subcomplex (Extended Data Fig. 1c). Acetylation of histone H3 at residue K14, a representative histone acetylation, was validated and the level of acetylation reached a plateau with increasing concentration of acetyltransferases, indicating nearly complete or highest level of acetylation in our experimental condition.

The purified PBAF was incubated with the acetylated nucleosome (nucleosome or NCP hereafter) in 1:1 stoichiometry in the presence of ATP analogue ADP-BeF_3_, followed by gradient fixation (Grafix) and cryo-EM sample preparation. Cryo-EM structure of PBAF-NCP was determined to overall resolution of approximately 4.4 Å (Extended Data Figs. 2, 3, Extended Data Table 1, Supplementary Videos 1, 2). The PBAF-NCP complex is organized into three modules including the Core, the actin-related proteins (ARP) (Cairns et al., 1998), the multi-subunit Base modules. Cryo-EM maps of the Base and Core modules were improved by focus refinement to near-atomic (3.4 Å to 4.1 Å) resolution and the ARP to 5.4 Å resolution. Structural models were built according to the cryo-EM maps with structure of BAF as reference (He et al., 2020) and aided by cross-linking mass spectrometry (XL-MS) analysis (Extended Data Fig. 4, Extended Data Table 2).

PBAF-NCP structure reveals a tripartite modular organization and is generally similar to the structures of nucleosome-bound human BAF (He et al., 2020; Mashtalir et al., 2020) and yeast SWI/SNF and RSC complexes (Han et al., 2020; Patel et al., 2019; Wagner et al., 2020; Ye et al., 2019) (Fig. 1b, Extended Data Fig. 5). SMARCA4 serves as a central scaffold that involves formation of three modules. The Core module is formed by the nucleosome-bound ATPase, a regulatory Hub connecting ATPase and ARP, and a Snf2 ATP coupling (SnAC) domain (Sen et al., 2011; Sen et al., 2013) packing against the surface of histone octamer (Figs. 1 and 2). The ARP module is formed by ACTB-ACTL6A heterodimer and helicase-SANT-associated (HSA) helix of SMARCA4 that bridges the Core and Base. Cryo-EM map reveals an additional density within that ARP that was not observed in the 10-subunit BAF structure (He et al., 2020), suggesting that this region is derived from PBAF-specific subunit or BCL7A (not present in the 10-subunit BAF). A two-stranded β-sheet of BCL7A was placed according to cryo-EM map and XL-MS (Extended Data Figs. 3, 4). The Base module accounts for the majority of molecular mass and is formed by the preHSA region of SMARCA4 and nine auxiliary subunits (Figs. 1 and 3).

**Figure 2.**
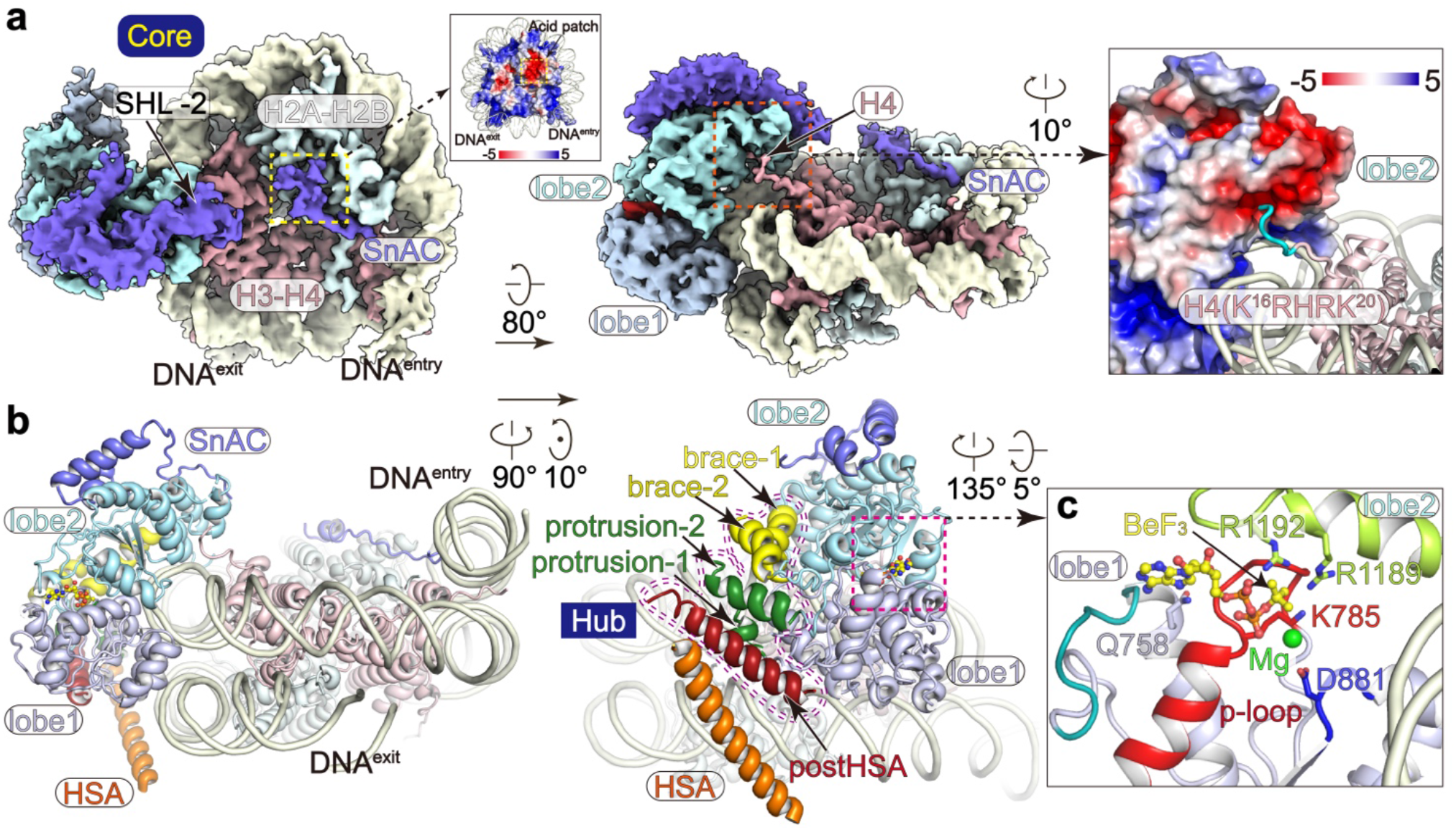
Structure of the nucleosome-bound ATPase. **(a)** Cryo-EM map of the Core module in two different views. Binding of SnAC to histone acidic surface is highlighted and binding of histone H4 tail to the acidic surface of ATPase lobe2 is shown in close-up view. (**b**) Structural model of the Core module with the Hub helices indicated. (**c**) Close-up view of the nucleotide-binding pocket with ADP-BeF_3_ and critical residues shown in sticks.

**Figure 3.**
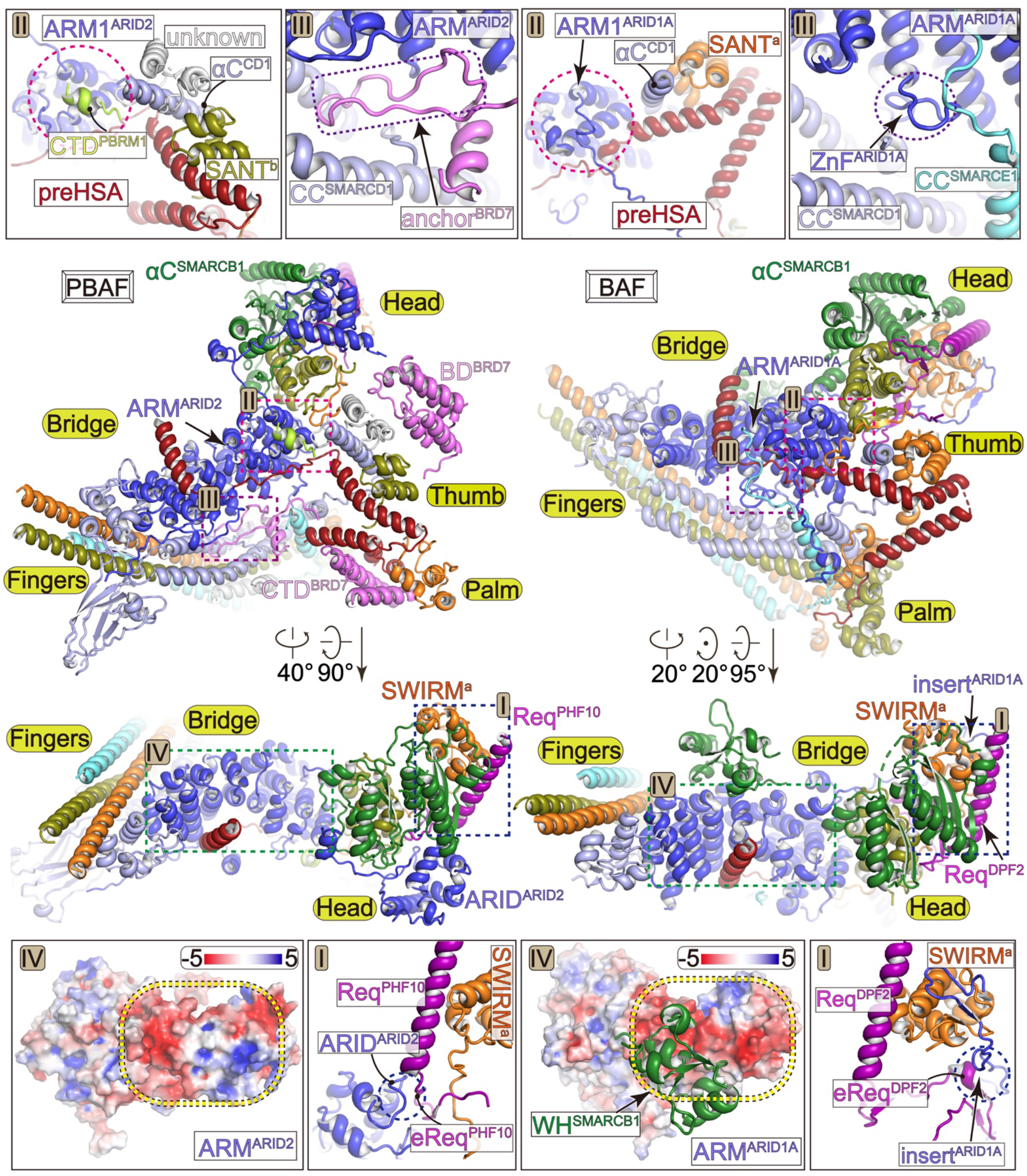
PBAF-specific organization of the Base module. Comparison of the Base modules of PBAF (left panels) and BAF (right panels) in two different views. Structural differences at region-I to region-IV are indicated on overall structure and shown in close-up views. The differences at equivalent positions are highlighted with dashed circles.

PBAF makes multiple contacts with nucleosome. Within the Core module, the ATPase stably grasps nucleosomal DNA at superhelical location (SHL) -2 and the interaction is buttressed by two tethers between ATPase-SnAC and histones (Figs. 1 and 2). Within the Base module, multiple DNA-binding and histone-binding domains/modules directly bind or are positioned near the nucleosome core particle (Figs. 1 and 4).

**Figure 4.**
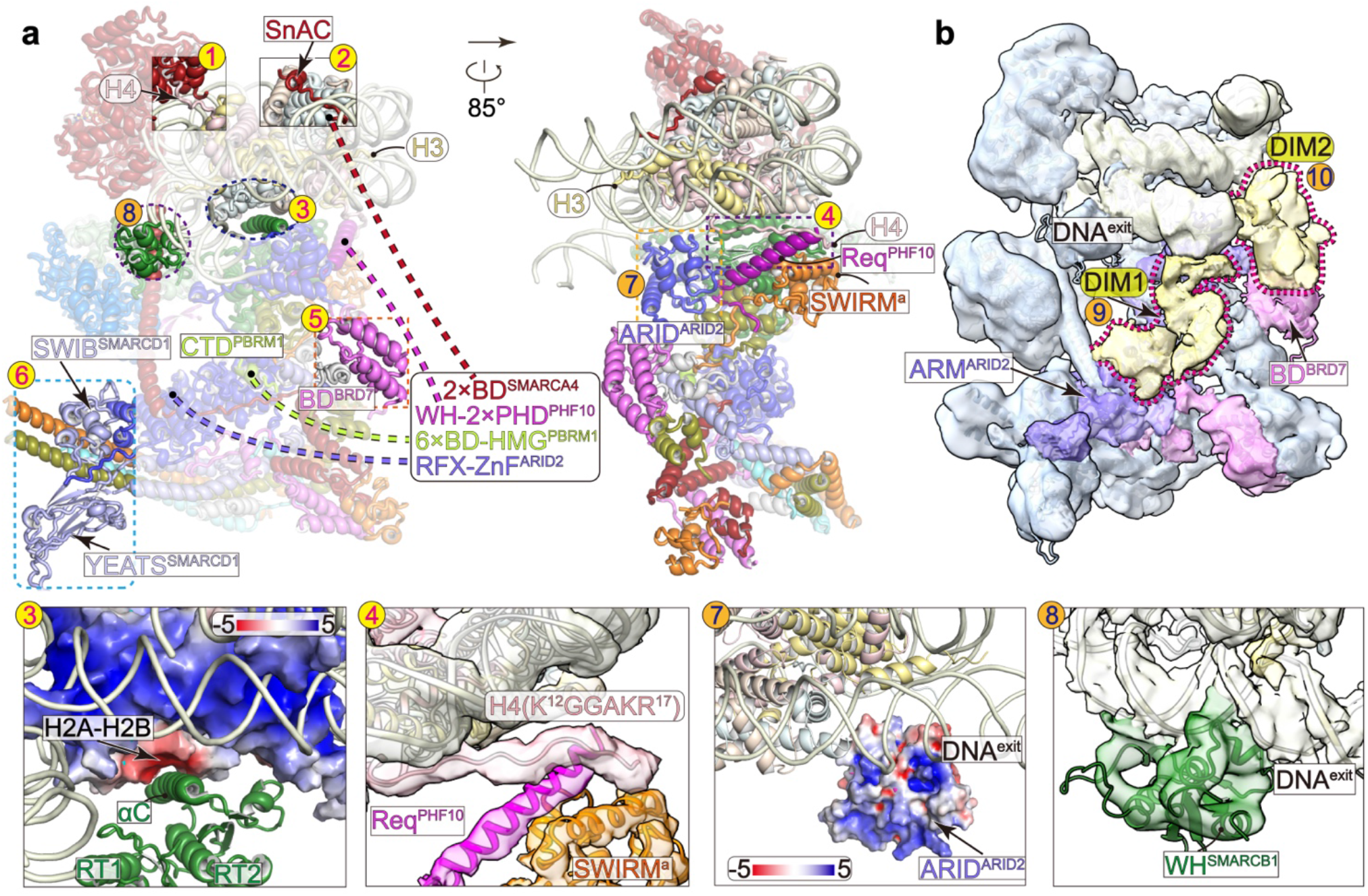
Histone/DNA-binding domains/modules. Structural model (**a**) and cryo-EM map at low threshold (**b**) of PBAF-NCP with the positions of histone/DNA-binding domains/modules indicated with numbers. Numbers 1 to 6 in yellow balls indicate the histone-binding domains and 7 to 10 in orange balls indicate the DNA-binding domains/modules. Invisible domains are connected with dashed lines to indicate their tethers with ordered regions. The bottom panels show close-up views of indicated contacts with involved regions shown in transparent cryo-EM map or electrostatic potential surface. Unassigned regions around exiting DNA, DIM1 and DIM2, are colored in yellowed and highlight with dashed lariats.

### The ATPase grasps nucleosomal DNA and the Hub couples the ATPase and Base

Structure of the ADP-BeF_3_-bound ATPase at near-atomic resolution shows network of interactions between the ATPase and ADP-BeF_3_, which results in a close conformation with the ATPase lobe1 and lobe2 grasping nucleosomal DNA (Fig. 2, Extended Data Fig. 6). As the mimetic γ-phosphate of ATP, BeF_3_ is stabilized by residues R1189 and R1192 of lobe2 and residue K785 within the P-loop of lobe1. The magnesium cation is coordinated by BeF_3_ and residue D881 of lobe1. Structural comparison of PBAF-NCP with nucleosome-bound ATPase of yeast Snf2 (PDB: 5Z3U) (Li et al., 2019) shows similar overall fold of the ATPase in ADP-BeF_3_-bound state, consistent with highly conserved catalytic mechanism. Compared to the isolated Snf2, the ATPase of PBAF slightly rotates and the Hub helices are displaced by up to 6 Å, likely resulted from the association of the Base through a regulatory Hub that connecting the HSA and ATPase.

The HSA-containing ARP and postHSA-containing Hub are essential for the function of SWI/SNF complexes and cancer-associated gain-of-function mutations are enriched on the Hub (Clapier et al., 2016; Clapier et al., 2020; Schubert et al., 2013; Szerlong et al., 2008). However, structural organization of the Hub was not fully understood. The PBAF-NCP structure shows that the Hub is formed by five α helices derived from postHSA, protrusion1/2, and brace1/2 (Fig. 2b, Extended Data Fig. 6). Two protrusion helices associate with ATPase lobe1 on one side and postHSA helix on the other side. Two brace helices form a helix hairpin, which associates with the two ATPase lobes and packs against the protrusion helices. Such domain organization suggests that the Hub couples the motions of ATPase and HSA-associated ARP-Base and therefore regulates chromatin remodeling activity.

### PBAF-specific modular organization of the Base

The Base of PBAF consists of six BAF/PBAF-shared subunits including SMARCA4, SMARCB1, SMARCD1, SMARCE1, and two SMARCC, and four PBAF-specific subunits including PBRM1, BRD7, ARID2 (counterpart of ARID1A/B in BAF), and PHF10 (counterpart of DPF1/2/3 in BAF) (Figs. 1, 3, Extended Data Fig. 7, Supplementary Videos 1 to 3). The Base module is divided into five submodules, including the Bridge that nucleates assembly of the Base, the Head that directly binds histone core particle, the Fingers formed by five-helix bundle and associated domains, the helical Thumb, and a split Palm. Consistent with the compositional similarity and differences, PBAF and BAF complexes share generally similar modular organization of the tripartite architecture. However, PBAF does exhibit PBAF-specific modular organization, distinct from that of BAF mainly at four regions, termed region-I to region-IV for simplicity.

### ARID2 serves as a scaffold for assembly of the Base

The central Bridge consists of a superhelical armadillo (ARM) repeat derived from the majority of ordered region in ARID2 (ARM^ARID2^), which covers residues 157-1817 and contains a structurally flexible insert (residues 480-1752) (Figs. 1, 3, Extended Data Fig. 7). Similar to that of BAF, the ARM^ARID2^ has seven ARM repeats (ARM1 to ARM7) arranged into a superhelical fold and serves as a rigid core to nucleate the Base formation through binding preHSA region of SMARCA4 and other Base subunits. ARM^ARID2^ and ARM^ARID1A^ exhibit compositional and conformational difference in ARM repeats and ARM-associated loops, generating distinct modular organization of PBAF and BAF.

Within the Head, the Req helix of PHF10 (Req^PHF10^) packs against a groove of SWIRM^a^ (one of the two SWIRM domains of two SMARCC) and repeat domain 2 (RPT2) of SMARCB1, similar to the binding pattern of Req^DPF2^ in BAF-NCP structure (He et al., 2020), consistent with highly conserved residues for the interaction (Fig. 3, Extended Data Figs. 8, 9). At the region-I, the N-terminal ARID domain of ARID2 (ARID^ARID2^) packs against the two RPT domains of SMARCB1 and an extension of Req domain (eReq) of PHF10. Specific incorporation of PHF10, instead of its counterpart DPF1/2/3, in PBAF may collectively result from the presence of eReq^PHF10^-binding ARID^ARID2^ and the lack of an insert of ARID1A, which stabilizes DPF1/2/3-specific eReq in BAF (Extended Data Figs. 7, 8d, 9).

### PBAF-specific PBRM1 and BRD7

Around the region-II of the Thumb, the C-terminal α helix of SMARCD1 (αC^SMARCD1^) is surrounded by a helix of preHSA of SMARCA4, a SANT domain of one SMARCC (SANT^b^), and three short α helices (Fig. 3, Extended Data Fig. 7). One of the three helices binds the Bridge on the ARM1^ARID2^ whereas this contact would generate steric clash with an ARM1^ARID1A^-associated loop in BAF. By contrast, the lack of PBRM1 in BAF leads to direct binding of the Thumb and Bridge and a rotation of the Thumb to the Head compared to that in PBAF. Cryo-EM map at near-atomic resolution and XL-MS analysis (Extended Data Figs. 3, 4) together support that this α helix is derived from the C-terminal domain of PBAF-specific PBRM1 (CTD^PBRM1^).

PBRM1 exclusively exists in PBAF and has no counterpart in BAF. Besides the CTD^PBRM1^, PBRM1 consists of six bromodomains (BD), a bromo-adjacent homology (BAH), and a high mobility group (HMG) domain that account for the majority of molecular mass but were not structurally observed in the cryo-EM map. A number of cancer-derived nonsense mutations resulted in truncations of PBRM1 that lack of CTD^PBRM1^ (Varela et al., 2011) (Extended Data Fig. 9b), consistent with the importance of CTD^PBRM1^ in assembly and function of PBAF complex.

Around the region-III, the Bridge and Fingers make direct contacts between ARM5-ARM7 and Fingers helix bundle (Fig. 3, Extended Data Figs. 7, 8). An anchor motif of BRD7 (anchor^BRD7^) inserts into the gap between the Bridge and Fingers and facilitates their interactions. By contrast, equivalent gap in BAF is occupied by two ARM^ARID1A^-associated loops coordinated by a zinc finger (He et al., 2020), which are absent in ARID2. The C-terminal domain of BRD7 (CTD^BRD7^) binds two N-terminal helices of SMARCA4 and a SANT domain of the other SMARCC (SANT^a^), generating a split Palm submodule that associated with the Thumb. The lack of CTD^BRD7^ in BAF allows this submodule to form a full Palm around the Fingers end through binding SMARCC helices and Pillar helices derived from SMARCD1 and SMARCE1.

Unexpectedly, the winged helix domain of SMARCB1 (WH^SMARCB1^) associates with ARM5^ARID1A^ of the Bridge in BAF (He et al., 2020) but was not observed in equivalent position in PBAF (Fig. 3, region-IV). By contrast, WH^SMARCB1^ in PBAF binds nucleosomal DNA at SHL +6.5 (Fig. 4, described below). Distinct positioning of WH^SMARCB1^ may result from the difference in charge distribution of WH^SMARCB1^-binding surface of ARM^ARID1A^ and equivalent region in ARM^ARID2^, which result from their sequence difference.

### The placements of multiple nucleosome-binding domains/modules

PBAF consists of over 20 nucleosome-binding domains/modules (Fig. 1a) that are thought to be functionally important in PBAF-mediated chromatin remodeling (Mashtalir et al., 2021). However, only a few of these domains/modules were observed in previously reported structures of SWI/SNF complexes (Han et al., 2020; He et al., 2020; Mashtalir et al., 2020; Patel et al., 2019; Wagner et al., 2020; Ye et al., 2019). Cryo-EM structure of PBAF-NCP reveals the placements of six histone-binding domains/motifs (Figs. 1, 4, Extended Data Fig. 8), including the H2A-H2B heterodimer-bound SnAC^SMARCA4^ and αC^SMARCB1^, the H4 tail-bound ATPase lobe2 and Req^PHF10^-SWIRM^SMARCC^ heterodimer, and histone-free BD^BRD7^ and YEATS-like domain of SMARCD1 (YEATS^SMARCD1^), and four DNA-binding domains/modules, including ARID^ARID2^ and WH^SMARCB1^, and two unassigned DNA-interaction modules (DIM1 and DIM2) around exiting DNA of the bound nucleosome.

### The Head and ATPase-SnAC bind nucleosome on two H2A-H2B heterodimers and two H4 tails

Locally refined cryo-EM map of the Core shows that the catalytic subunit SMARCA4 stably engages with nucleosome (Figs. 2a, 4a). Apart from DNA-ATPase interaction, the acidic patch of ATPase lobe2 binds the N-terminal positively charged tail (K^16^RHRK^20^) of histone H4 (Fig. 4a, position-1) and the SnAC domain extends out of the ATPase domain and winds over the acidic patch of H2A-H2B heterodimer (Fig. 4a, position-2). The SnAC extends to the nucleosomal DNA at SHL -6. The two nucleosome-SMARCA4 tethers may facilitate nucleosome-association of the ATPase during DNA translocation, consistent with the known functions of SnAC domain (Sen et al., 2011; Sen et al., 2013) and K16/20 acetylation of histone H4 (Mashtalir et al., 2021) in regulating activities of SWI/SNF complexes.

The positively charged helix αC^SMARCB1^ binds the acidic patch of the bottom H2A-H2B heterodimer and binding pattern is similar to that observed in BAF-NCP structure (He et al., 2020) (Fig. 4, position-3). In PBAF and BAF complexes, the Core and Base are similarly connected by the H2A-H2B heterodimer-αC^SMARCB1^ and the Hub-HSA-ARP tethers (Extended Data Fig. 7). Some cancer-associated mutations are enriched on αC^SMARCB1^ (loss of function) (Valencia et al., 2019) and the Hub (gain of function) (Clapier et al., 2020), supporting their functional importance in both BAF and PBAF complexes.

Due to the differences in composition and arrangement of nucleosome-binding domains, PBAF and BAF exhibit distinct intermodular organization of the Core and Base (Fig. 4, Extended Data Fig. 7, Supplementary Video 3). For example, compared to that of BAF, the Head of PBAF rotates toward nucleosomal DNA at +2, permitting the binding of the N-terminal tail of histone H4 to Req^PHF10^-SWIRM^SMARCC^ heterodimer (Fig. 4a, position-4, Extended Data Fig. 8b). The interaction is likely mediated by positively charged residues of H4 tail and a negatively charged patch of Req^PHF10^-SWIRM^SMARCC^ heterodimer.

### Positions of two putative histone-binding domains

We observed relatively weak cryo-EM map positioned near CTD^PBRM1^ within the Thumb (Fig. 4, position-5, Extended Data Fig. 2). XL-MS analysis suggests that it is derived from the bromodomain of BRD7 (BD^BRD7^) (Extended Data Fig. 4, Extended Data Table 2). Predicted structural model of BD^BRD7^ was placed into the map and the putative histone binding site is about 70 and 50 Å away from the histone fold regions of H3 and H4, respectively (Extended Data Fig. 8). Thus, BD^BRD7^ is accessible to acetylated histone tails, ∼30-40 residues in length, in line with its predicted function in binding of histone acetylation.

Within the Bridge, the SWIB domain of SMARCD1 is organized by four short helices and serves as a helical extension of the ARM repeat (Fig. 4, position-6, Extended Data Fig. 8a). The YEATS-like domain of SMARCD1 (YEATS^SMARCD1^) associates with SWIB^SMARCD1^ and the coiled-coil of SMARCD1 in the Fingers. YEATS^SMARCD1^ is organized into two parallel four-stranded β-sheets, as in other YEATS domains, but lacks a conserved acetyl-lysine binding pocket. As a peripheral domain, YEATS^SMARCD1^ is positioned by as far as 120 Å away from nucleosome core particle, suggesting its role independent of binding nucleosome. Nevertheless, the YEATS-like domain is strictly conserved among the snf12/Rsc6/SMARCD1 family proteins, which regulates expression of genes in stress response (Cairns et al., 1996; He et al., 2021).

### DNA-binding domains around the exiting DNA

Apart from the ATPase, additional DNA-binding domains were observed around the exiting DNA (Fig. 4, position-7). ARID domains of ARID1A/1B and ARID2 are predicted to bind DNA (Patsialou et al., 2005) but not structurally observed in previous studies of SWI/SNF complexes. ARID^ARID2^ in the Head submodule has no direct contact with nucleosomal DNA but its positively charged surface faces toward DNA, leaving the possibility to make a direct DNA-binding if underwent conformational change.

Cryo-EM map of PBAF-NCP shows noticeable density associated with nucleosomal DNA at SHL +6.5 and structural model of WH^SMARCB1^ fits this density well (Fig. 4, position-8, Extended Data Fig. 8c). The placement of WH^SMARCB1^ suggests that the positively charged helix insets into the major groove of DNA with a cluster of characteristic basic residues (R37/R40/K45/R46/R52/R53) positioned near phosphate groups for charge-charge interactions, exhibiting a DNA-binding pattern similar to previously proposed model (Allen et al., 2015).

Cryo-EM map reveals an unassigned density of DIM1 connecting ARM helices of the Bridge and extranucleosomal DNA at 10-bp to 20-bp downstream of the exit site at SHL +7, suggesting that DIM1 is possibly derived from the RFX-like DNA binding domain (RFX) and/or zinc finger (ZnF) of ARID2 (Fig. 4b, position-9, Extended Data Fig. 8e). Another unassigned region (DIM2) associates with the BD^BRD7^ and makes contacts with extranucleosomal DNA (Fig. 4b, position-10, Extended Data Fig. 8e). This region is possibly derived from DNA-binding domains of nearby subunits, such as WH domain of PHF10, HMG domain of PBRM1, and C2H2 zinc finger of ARID2.

PBAF-DNA contacts are enriched around the exiting site of nucleosomal DNA, a characteristic feature distinct from that of BAF-NCP structure, in which neither equivalent DNA contact nor placement of DNA-binding domains was observed (He et al., 2020). Cryo-EM map of nucleosome-bound RSC (yeast counterpart of PBAF) at low resolution showed that an unassigned DIM binds extranucleosomal DNA 20-bp to 40-bp downstream of SHL +7 of nucleosome (Wagner et al., 2020) (Extended Data Fig. 8f). While DNA-binding pattern differs in PBAF and RSC, binding of exiting DNA may represent an evolutionarily conserved function in targeting PBAF/RSC to promoters (Badis et al., 2008; Kubik et al., 2015; Kubik et al., 2018; Lorch et al., 2014).

Apart from the observed histone-binding and DNA-binding domains/modules, PBAF also consists of domains/modules that are invisible in the cryo-EM map, including two bromodomains of SMARCA4, WH domain and two PHD domains of PHF10, and HMG and six bromodomains of PBRM1, which are tethered with the Core module, the Head and Thumb submodules, respectively, consistent with their predicted roles in binding DNA and modified histone tails and targeting chromatin for function (Fig. 4).

## Discussion

PBAF and BAF share identical subunits and equivalent subunits, and therefore are commonly considered highly similar in structure and function. PBAF differs from BAF in its presence of multiple acetylation-binding bromodomains. However, our study unexpectedly shows marked structural difference in their modular organization of the Base and placements of DNA/histone-binding domains. PBAF-specific subunits, ARID2, PBRM1, PHF10, and BRD7 not only provide PBAF-specific nucleosome-binding domains but also alter the placements of some nucleosome-binding domains of the PBAF/BAF-shared subunits, such as WH^SMARCB1^ and Req^PHF10^-SWIRM^SMARCC^. Consistent with their functional requirement, the majority of nucleosome-binding domains exist or associate with the Head and Thumb submodules, which are positioned near the nucleosome. Such multivalent nucleosome-binding pattern was not observed in previous studies. The PBAF-NCP structure may provide a framework to further investigate whether, and if yes, how these nucleosome-binding domains work coordinately or redundantly in integrating chromatin marks for remodeling chromatin targets.

Cancer-derived mutations are also frequently observed in PBAF-specific subunits (Hakimi et al., 2020; Varela et al., 2011) (Extended Data Fig. 9b). Missense mutations predominately occur on domains that are involved in complex assembly (binding of other subunits) or histone/DNA-binding domains, in line with the critical roles of these domains in PBAF function. Large number of nonsense mutations occur throughout PBRM1 and ARID2, consistent with the importance of CTD^PBRM1^ and ARM7^ARID2^ in organizing the Base module.

During preparation of this manuscript, a structure of human PBAF-NCP was published (Yuan et al., 2022). The complex was assembled with unmodified nucleosome and 12-subunit PBAF (lack of BCL7A). To improve complex behavior in structure determination, Yuan et al. removed the N-terminal region of SMARCA4 (residues 1-159), the first four bromodomains of PBRM1 (residues 1-630), and the internal region of ARID2 (residues 627-1591). Cryo-EM map at high resolution indeed favor the assignment of Base subunits, including CTD^PRMB1^ and BD^BRD7^. By contrast, we assembled the complex using acetylated nucleosome and human PBAF containing 13 full-length subunits. The generated cryo-EM map revealed subunits BCL7A and YEATS^SMARCD1^ domain and additional PBAF-NCP contacts including DNA-WH^SMARCB1^, DNA-DIM1, DNA-DIM2, and histone H4 tail-Req^PHF10^-SWIRM^SMARCC^. Thus, our study provides complementary structural insights into multivalent interactions between PBAF and nucleosome and a framework for understanding PBAF functions in chromatin targeting and remodeling.

## Acknowledgments

We thank the Center of Cryo-Electron Microscopy of Fudan University for the supports on data collection and the Biomedical Core Facility, Fudan University for the support on mass spectrometry analyses. This work was supported by grants from the National key R&D program of China (2016YFA0500700), the National Natural Science Foundation of China (32030055, 31830107, 31821002, 31970146), the National Postdoctoral Program for Innovative Talent (S. H.), the Shanghai Municipal Science and Technology Major Project (2017SHZDZX01), Shanghai Municipal Science and Technology Commission (19JC1411500), the National Ten-Thousand Talent Program (Y. X.), the National Program for support of Top-Notch Young Professionals (Y. X.), and the XPLORER Prize (Y. X).

## Author contributions

L. W. prepared the samples for structural and biochemical analyses with help from J. Y. and S. H.; Z. Y. performed EM analyses and model building with help from Q. W. and Z. C.; Y. X., L. W. and S. H. wrote the manuscript; Y. X. supervised the project.

## Data and materials availability

Cryo-EM maps and atomic coordinates will be deposited in the EMDB and PDB upon the acceptance of this manuscript.

## Methods

### PBAF expression and purification

The thirteen full-length open reading frames (ORFs) of PBAF subunits were subcloned into modified pMLink vector containing no tag or the N-terminal Flag tag and 4 x Protein A tag followed by an HRV-3C cleavage site. All human PBAF subunits were co-transfected into suspension Expi293F cells using polyethylenimine (Polysciences). Cells were cultured for 72 h at 37°C and harvested by centrifugation. For complex purification, all the steps were performed at 4°C. Cells were disrupted in lysis buffer containing 50 mM HEPES pH 8.0, 300 mM NaCl, 5% (v/v) Glycerol, 0.2% (w/v) chaps, 2 mM MgCl2, 0.5 mM EDTA (Ethylenediaminetetraacetic Acid), 2 mM DTT (Dithiothreitol), 1 mM PMSF (Phenylmethylsulfonyl fluoride), 1 μg/mL Aprotinin, 1 μg/mL Pepstatin, 1 μg/mL Leupeptin for 30 min. Cell lysate was clarified by centrifugation at 16000 rpm for 30 min. The supernatant was incubated with IgG resin for 4 h and washed thoroughly with wash buffer containing 20 mM HEPES pH 8.0, 150 mM KCl, 5% (v/v) Glycerol, 0.1% chaps, 2 mM MgCl2, 2 mM DTT. After on-column digestion overnight, immobilized protein was eluted using wash buffer and further loaded onto an ion-exchange column (MonoQ 5/50 GL column, GE Healthcare) to achieve highly pure PBAF complex. The peak fractions corresponding to PBAF complex were collected and concentrated to ∼3 mg/mL. The concentrated samples were used for subsequent biochemical and structural analyses.

### Preparation of nucleosomes

Canonical human histone H2A-H2B heterodimer and H3.1-H4 heterotetramer were separately co-expressed as soluble protein in Escherichia coli BL21 (DE3) cells as described previously (Klinker et al., 2014). In brief, cells were disrupted in lysis buffer containing 50 mM Tris-HCl (pH 8.0), 2 M NaCl, 5% (v/v) glycerol, 0.7 mM β-mercaptoethanol (β-ME), and then purified through ion-exchange chromatography. For histone octamer assembly, H2A-H2B heterodimer in 1.2-fold excess was mixed with H3.1-H4 heterotetramer and then incubated for 0.5 h at 4°C, followed by a size exclusion chromatography (Superdex 200 10/300, GE Healthcare). Peak fractions were concentrated and used for nucleosome assembly.

DNA fragments for mononucleosome reconstitution were prepared by PCR amplification (Maskell et al., 2015). Three different mononucleosome DNA used in this study contained the Widom 601 positioning sequence (Lowary and Widom, 1998). Nucleosome 45N45 (N denotes nucleosome) consists of two flanking sequences of 45-bp in length. Nucleosome 0N90 consists of one flanking DNA with 90-bp in length. Nucleosome 15N51 consists of a 15-bp and a 51-bp flanking DNAs. nA center-positioned nucleosome 45N45 was assembled and used as a substrate in nucleosome remodeling activity. The DNA sequence of 45N45 is as below:

GCATCCCTTATGTGAGGTACCCTATACGCGGCCGCCCCGGATCCCCTGGAGAATCCCGGTGCCGAGGCCGCTCAATTGGTCGTAGACAGCTCTAGCACCGCTTAAACGCACGTACGCGCTGTCCCCCGCGTTTTAACCGCCAAGGGGATTACTCCCTAGTCTCCAGGCACGTGTCACATATATACATCCTGTTCCAGTGCCGGGCATGTATTGAACAGCGTTTAAACCGGTGCCAGT(the ‘601’ positioning sequence is underscored).

An end-positioned nucleosome 0N90 was assembled and used as the reference in nucleosome remodeling activity. The DNA sequence of 0N90 is as below:

CTGGAGAATCCCGGTGCCGAGGCCGCTCAATTGGTCGTAGACAGCTCTAGCACCGCTTAA ACGCACGTACGCGCTGTCCCCCGCGTTTTAACCGCCAAGGGGATTACTCCCTAGTCTCCAGGCACGTGTCACATATATACATCCTGTTCCAGTGCCGGTGTCGCTTGGGTCCCGAGGTATTCAAGCTTATCGATACCTGCGACCACGAGGGGGGGTGCCGGGCATGTATTGAACAGC(the ‘601’ positioning sequence is underscored).

The nucleosome 15N51 was assembled and used for cryo-EM. The DNA sequence of 15N51 is as below:

ATCCTGGGGAATTCCCTGGAGAATCCCGGTGCCGAGGCCGCTCAATTGGTCGTAGACAGCTCTAGCACCGCTTAAACGCACGTACGCGCTGTCCCCCGCGTTTTAACCGCCAAGGGGATTACTCCCTAGTCTCCAGGCACGTGTCACATATATACATCCTGTTCCAGTGCCGGTGTCGCTTGGGTCCCGAGGATTACAAGCTTATCGATAGAT(the ‘601’ positioning sequence is underscored).

The 15N51 DNA fragment was inserted into the pUC57 vector, and the plasmid DNA was amplified in the *E. coli* strain DH5α. The 15N51 DNA fragment was excised from the plasmid DNA by EcoR V. The 45N45 and 0N90 DNA fragments were prepared using PCR amplification. All the nucleosomal DNA fragments were purified using ion-exchange chromatography (Source Q 5/5, GE Healthcare) and isopropanol precipitation. The purified DNA pellet was dissolved in 1 x TE buffer.

Nucleosome reconstitution was performed by mixing DNA with octamer at a equimolar ratio, with a linear salt gradient dialysis according to previously published research (Luger et al., 1999). Finally, nucleosomes were dialyzed to 1 x HE buffer (10 mM HEPES pH 8.0, 0.1 mM EDTA). The nucleosomes can be immediately used for complex assembly and biochemical assay.

### In vitro chromatin remodeling assay

In vitro chromatin remodeling assay was performed using purified PBAF complex and nucleosome 45N45 as substrate. Nucleosome 45N45 (120 nM) was mixed with PBAF (80 nM) in buffer containing 20 mM Tris-HCl (pH 8.0), 60 mM KCl, 5 mM MgCl2, 0.1 mg/mL BSA, 5% (v/v) glycerol. The reactions were started with the addition of 1 mM ATP at 30°C and stopped at different time points (0, 0.25, 0.5, 3, 5, 10, 30 min) by adding competitor plasmid (∼1 ug) in excess, followed by further incubation for 30 min on ice. The reaction samples were analyzed by 5% native PAGE gel at 4°C and run in 0.5 x Tris-Glycine buffer for 50 min at 180V constant. The PAGE gels were stained with GelRed dye and visualized using the Tanon-2500 image system. Ejection of H2A-H2B heterodimer as validated by western blotting using antibody against histone H2B.

### Nucleosome acetylation

To generate acetylated nucleosome, we purified two histone acetyltransferases (HATs), p300 and SAGA acetyltransferase subcomplex containing KAT2A, STAF36, TADA2B, and TADA3L. The expression and purification procedures are essentially similar to that of PBAF. Nucleosome 15N51 was acetylated in reaction buffer containing 50 mM Tris-HCl (pH 8.0), 0.1 mM EDTA, 10% (v/v) glycerol, 1 mM DTT, 1 mM PMSF. Nucleosome (1.6 μM) was first incubated with increasing concentration of the mixture of HATs at 30°C for 5 min, followed by the addition of 50 μM acetyl-CoA for another 30 min at 30°C (Chatterjee et al., 2011). Western blotting was used to detect the acetylation efficiency of nucleosome. For large-scale preparation of the acetylated nucleosome, 15N51 nucleosome and HATs were incubated in 1:1 stoichiometry with the reaction performed as mentioned above. The two HATs were separated from the PBAF-NCP complex during Grafix.

### Complex assembly and gradient fixation

The PBAF-NCP complex was cross-linked and purified using gradient fixation (Grafix) (Kastner et al., 2008). In brief, the purified PBAF and acetylated nucleosome 15N51 were mixed at a ratio of 1:1 for 30 min at 4°C followed by the addition of 2 mM MgCl_2_, 0.5 mM ADP, 7 mM NaF, 1 mM BeSO_4_ and incubation by 15 min at 30°C. The assembled sample was loaded onto a gradient generated from a glycerol light solution containing 15% (v/v) glycerol, 20 mM HEPES pH 7.0, 60 mM KCl, 2 mM MgCl_2_, 0.5 mM ADP, 7 mM NaF, 1 mM BeSO_4_, 2 mM DTT and a glycerol heavy solution containing 35% (v/v) glycerol, 20 mM HEPES pH 7.0, 60 mM KCl, 2 mM MgCl_2_, 0.5 mM ADP, 7 mM NaF, 1 mM BeSO_4_, 2 mM DTT and 0.01% (v/v) glutaraldehyde. The sample was subjected to ultracentrifugation at 38000 rpm for 15h in an SW41Ti swinging-bucket rotor (Beckman) at 4 °C. The peak fractions of the cross-linked PBAF-NCP complex was concentrated and dialyzed overnight against a buffer containing 20 mM HEPES pH 7.0, 60 mM KCl, 2 mM MgCl_2_, 0.5 mM ADP, 7 mM NaF, 1 mM BeSO_4_, 2 mM DTT, followed by cryo-EM grids preparation.

### Cryo-EM sample preparation

For negative staining EM grid preparation, samples (5 μL at a concentration of ∼0.06 mg/mL) were applied onto glow-discharged copper grids supported by a continuous thin layer of carbon film for 60 s before negative staining by 2% (w/v) Uranyl Acetate solution at room temperature. The grids were prepared in the Ar/O_2_ mixture for 15 s using a Gatan 950 Solarus plasma cleaning system with a power of 35 W. The negatively stained grids were loaded onto a Thermo Fisher Scientific Talos L120C microscope equipped with a Ceta CCD camera and operated at 120 kV at a nominal magnification of 92,000 x, corresponding to a pixel size of 1.58 Å on the specimen.

For cryo-EM grid preparation, samples (4 μL at a concentration of ∼0.6 mg/mL) were applied to freshly glow-discharged Quantifoil R1.2/1.3 holey carbon grids. After incubation for 5 s at 6 °C and 100% humidity, the grids were blotted for 1 s with blot force 2 in a Thermo Fisher Scientific Vitrobot Mark IV and plunge-frozen in liquid ethane at liquid nitrogen temperature. The grids were prepared in the H_2_/O_2_ mixture for 60 s using a Gatan 950 Solarus plasma cleaning system with a power of 5 W. The ø 55/20 mm blotting paper is made by TED PELLA used for plunge freezing.

### Cryo-EM data collection

The cryo-EM grids were loaded onto a Thermo Fisher Scientific Titan Krios transmission electron microscope operated at 300 kV for data collection. Cryo-EM images were automatically recorded by a post-GIF Gatan K3 Summit direct electron detector in the super-resolution counting mode using Serial-EM with a nominal magnification of 64,000 x in the EFTEM mode, which yielded a super-resolution pixel size of 0.667 Å on the image plane, and with defocus values ranging from -1.0 to -2.5 μm. Each micrograph stack was dose-fractionated to 40 frames with a total electron dose of ∼50 e^−^/Å^2^ and a total exposure time of 3.6 s. 9,156 micrographs of PBAF-NCP were collected for further processing.

### Image processing

Movie stacks were corrected for drift and beam-induced motion correction by MotionCor2 (Zheng et al., 2017) with binned 2-fold to a calibrated pixel size of 1.334 Å/pixel, which generated drift-corrected summed micrographs with and without electron-dose weighting. The defocus values were estimated by Gctf (Zhang, 2016) from non-dose-weighted summed images. Other procedures of cryo-EM data processing were performed within RELION v3.0 (Kimanius et al., 2016; Scheres, 2012) and cryoSPARC v3 (Punjani et al., 2017) using the dose-weighted micrographs.

Particles were automatically picked and subjected to reference-free 2D classification, yielding a total of 2,071,055 particles. The particles were further subjected to the 3D classifications. The classes with good quality consisted of 254,834 particles and were subsequently subjected to the 3D classification with Base module mask. And 92,526 particles were selected from good 3D classes, which were used for 3D classification in cryoSPARC v3. A final set of 37,528 homogeneous PBAF-NCP complex particles were selected to perform a final 3D reconstruction in cryoSPARC, yielding a reconstruction of PBAF-NCP complex at 4.4 Å resolution. Local refinement focused on the Core module with mask could reconstitute the Core module at 3.4 Å resolution. To improve the map quality of the Base module and the ARP module, the signal of Core module was subtracted from two classes of 3D classification containing 81,287 particles. The subtracted particles were further subjected to 2D classification, yielding 76,554 particles after clearance. A further 3D classification by applying mask for the ARP module and the Base module resulted in a clean dataset containing 31,995 particles. The resulting particles were refinement in cryoSPARC, yielding a reconstruction of ARP and Base module at 4.2 Å resolution. Local refinement focused on the ARP module and the Base module with mask generated reconstructions of the ARP module and the Base module at 5.4 Å and 4.1 Å, respectively.

All reported resolutions are based on the gold-standard Fourier shell correlation (FSC) = 0.143 criterion. The GSFSC curves were corrected for the effects of a soft mask with high-resolution noise substitution. All cryo-EM maps were sharpened by applying a negative B-factor estimation in cryoSPARC Sharpening Tools. All the visualization and evaluation of the 3D volume map were performed within UCSF Chimera or UCSF ChimeraX (Pettersen et al., 2004), and the local resolution variations were calculated using cryoSPARC.

### Model building and structure refinement

The structures of RSC complex (PDB: 6TDA) (Eustermann et al., 2018; Wagner et al., 2020; Ye et al., 2019), (PDB:6KW4) (Ye et al., 2019) and structure of BAF complex (PDB:6LTJ) (He et al., 2020) were used as initial structural templates, which were fitted into the cryo-EM maps by rigid-body fitting using UCSF Chimera followed by iterative rounds of manual adjustment and rebuilding in COOT (Emsley and Cowtan, 2004). The model was finalized by rebuilding in ISOLDE (Croll, 2018) followed by refinement in Phenix (Adams et al., 2002) with secondary structure and geometry restraints using the cryo-EM maps. Overfitting of the model was monitored by refining the model in one of the two half maps from the gold-standard refinement approach and testing the refined model against the other map. Statistics of the map reconstruction and model refinement can be found in Extended Data Table 1. The final structural model was validated using Phenix. Map and model representations in the figures and movies were prepared by PyMOL (https://pymol.org/), UCSF Chimera or UCSF ChimeraX.

### Cross-linking and mass spectrometry analysis

The Cross-linking Mass Spectrometry (XL-MS) analysis was performed as previously described (Chen et al., 2021). The purified PBAF complex (0.45 μM) was incubated with nucleosome at a ratio of 1:1 in the presence of ADP-BeF_3_ followed by cross-linking MS analyses. The PBAF-NCP complex was incubated with DSS (250 μM) at 25°C with shaking at 500 rpm (ThermoMixer) for 1 h. Reaction was terminated by adding 20 mM ammonium bicarbonate (Sigma). The cross-linked sample was precipitated with cooled acetone and dried in a speed vac. The pellet was dissolved in 8 M Urea, 100 mM Tris-HCl pH 8.5, followed by TCEP reduction, iodoacetamide (Sigma) alkylation, and trypsin (Promega) digestion overnight at 37°C using a protein/enzyme ratio of 50:1 (w/w). Tryptic peptides were desalted with Pierce C18 spin column (GL Sciences) and separated in a proxeon EASY-nLC liquid chromatography system by applying a step-wise gradient of 0-85% acetonitrile (ACN) in 0.1% foricacid. Peptides eluted from the liquid chromatography were directly electrosprayed into the mass spectrometer with a distal 2 kV spray voltage. Data-dependent tandem mass spectrometry (MS/MS) analyses were performed on Thermo Q-Exactive instrument in a 60-minute gradient. The acquired raw data files were processed with pLink2 software (Chen et al., 2019) and the results were visualized using the xiNET online server (Combe et al., 2015).

**Extended Data Figure 1.**
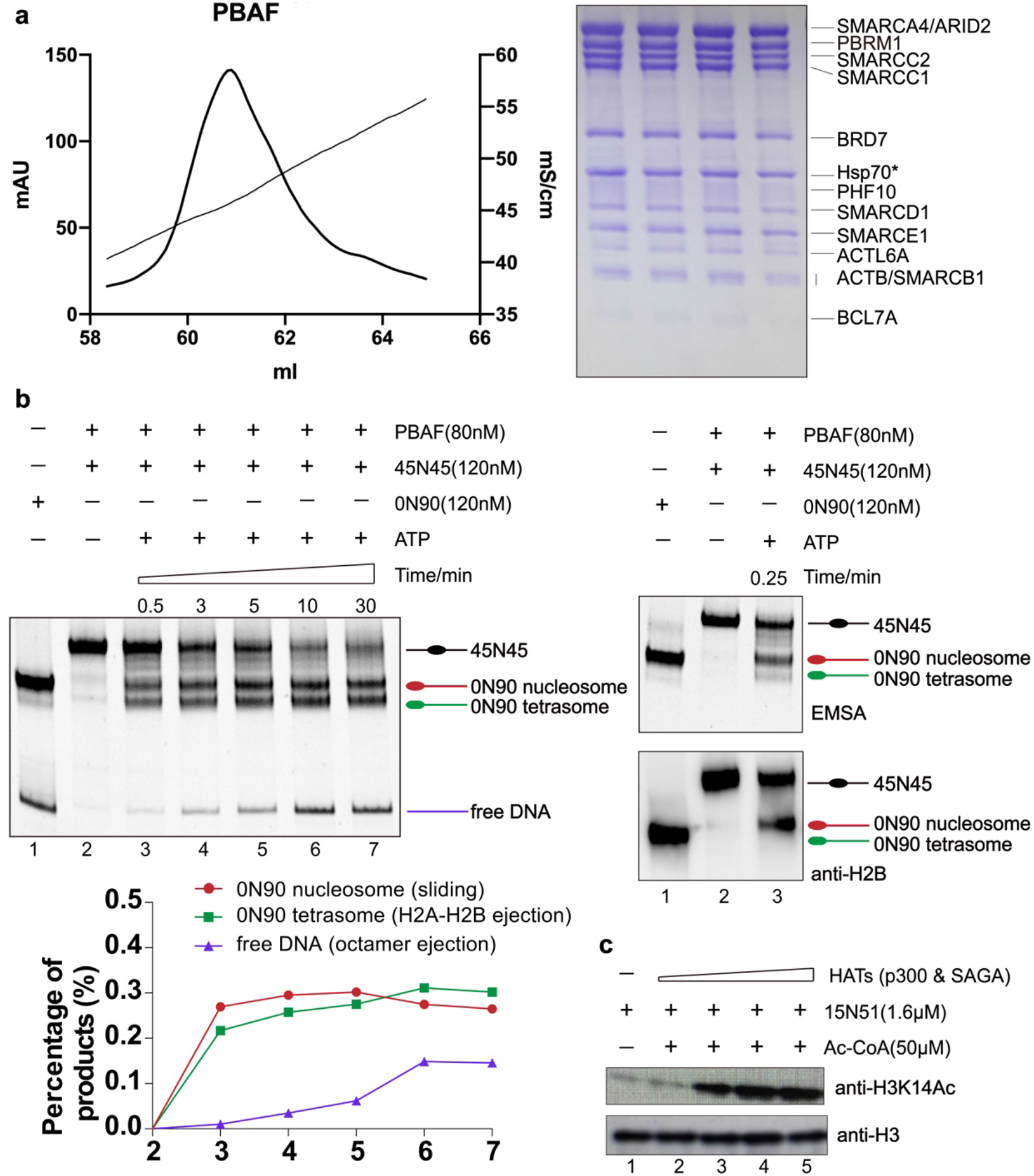
Protein purification and remodeling activities of PBAF complex. **(a)** Purification of PBAF complex. Profile of ion-exchange purification of the 13-subunit PBAF complex. Peak fractions were subjected to SDS-PAGE followed by Coomassie blue staining. (**b**) In vitro chromatin remodeling assay shows nucleosome sliding and ejection activities of PBAF. Reconstituted nucleosome 0N90 serves as a reference for an end-positioned nucleosome, the product of chromatin sliding reaction. The generated 0N90 tetrasome represents the product of H2A-H2B ejection with the ejection of H2A-H2B dimer confirmed using antibody against histone H2B. Free DNA represents the product of histone octamer ejection. (**c**) In vitro acetylation of nucleosome by increasing concentration of a mixture of two acetyltransferases, p300 and SAGA acetyltransferase subcomplex. The level of acetylation was detected using antibody against acetylated histone H3K14.

**Extended Data Figure 2.**
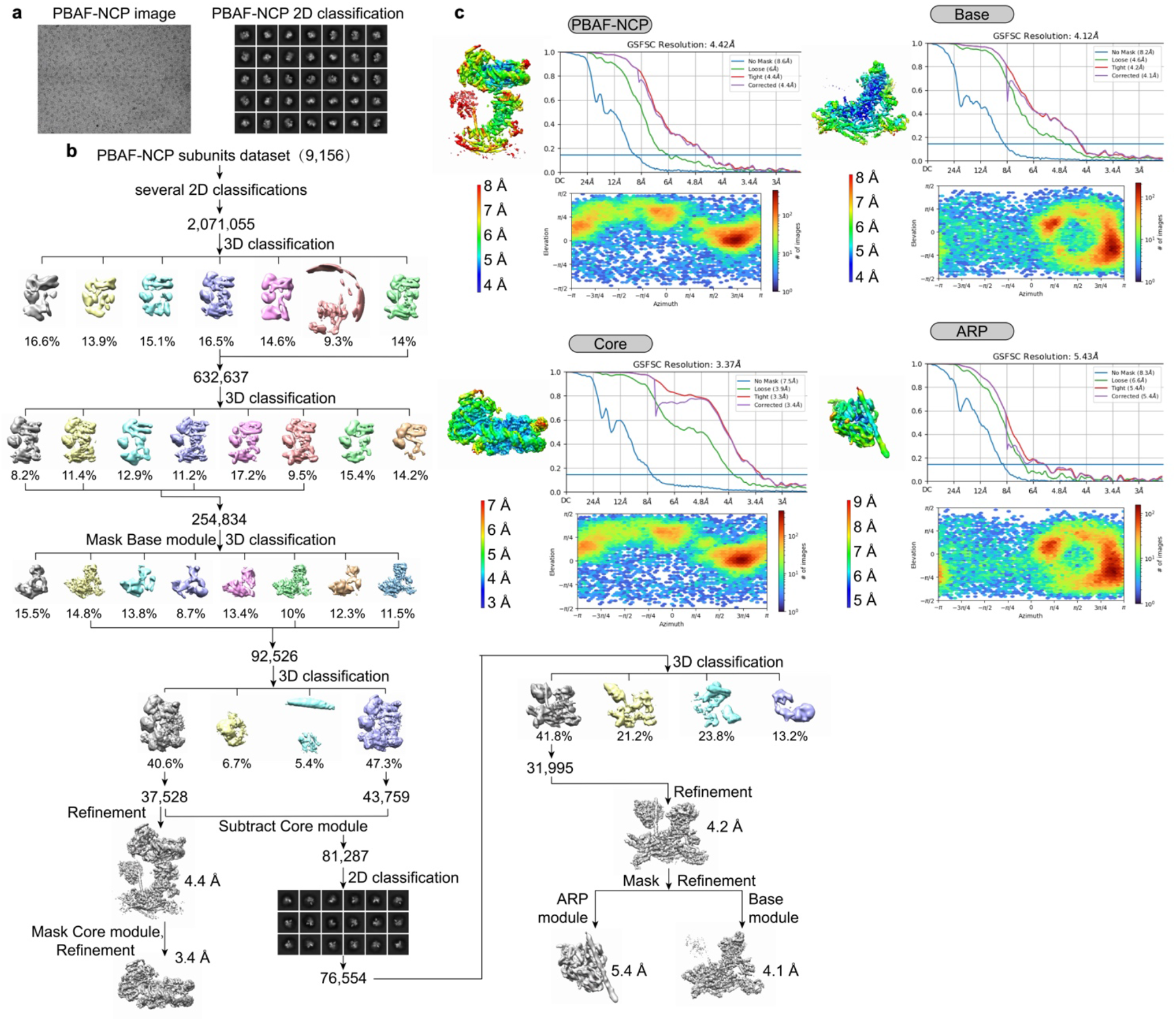
Data collection and image processing of PBAF-NCP complexes. (**a-b**) Representative cryo-EM images, 2D classification (b) and flow-charts of the cryo-EM image processing (b) of PBAF-NCP sample. (**c**) Local resolution estimation, GSFSC curves and direction distribution of the cryo-EM reconstructions of whole complex and the Core, Base and ARP modules.

**Extended Data Figure 3.**
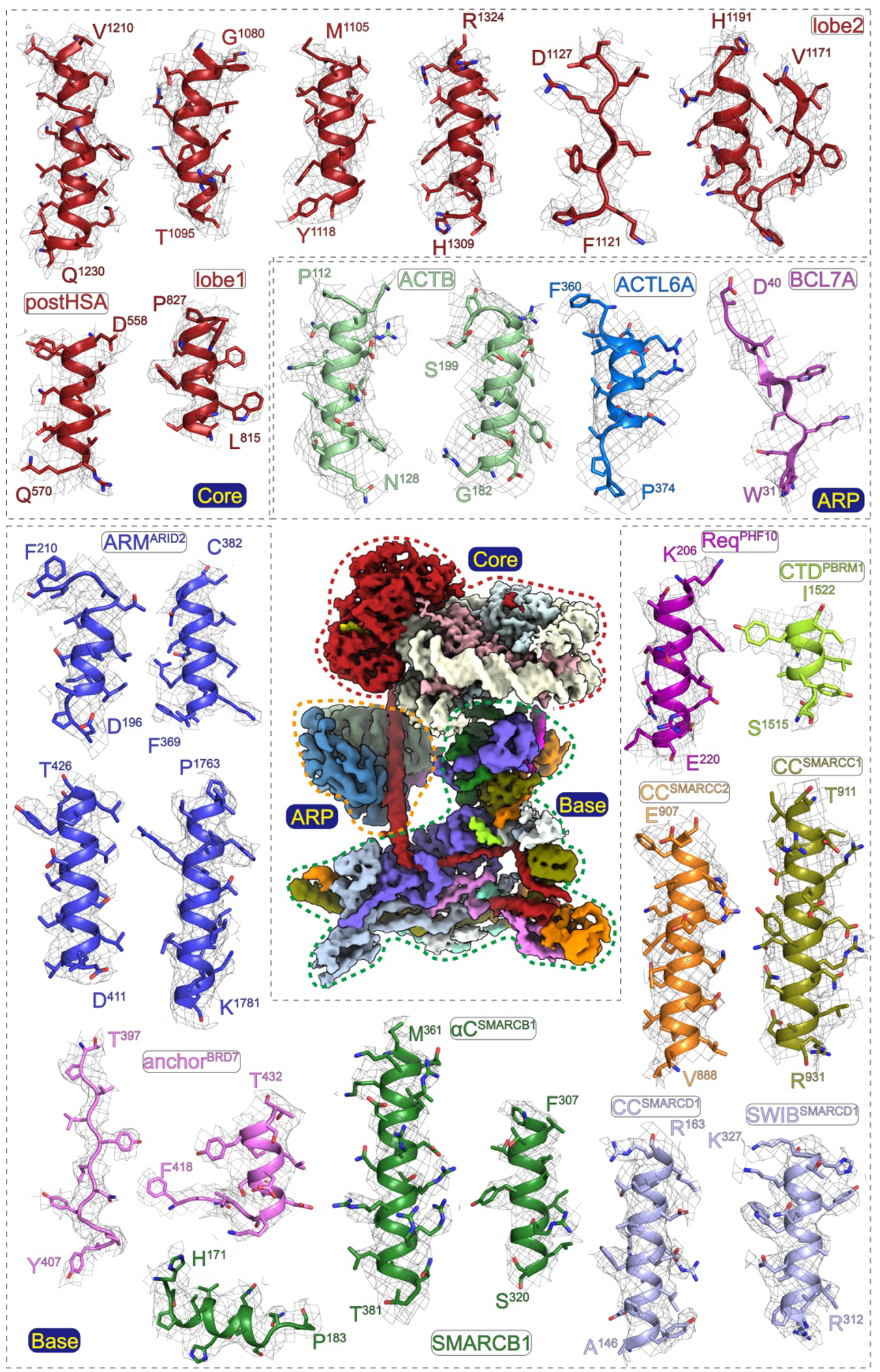
Cryo-EM map and structural model. Composite cryo-EM map of the Core module (3.4 Å), Base module (4.1 Å), and ARP module (5.4 Å). Representative regions of PBAF subunits are shown in close-up views. Structural models shown in sticks representation are well covered by cryo-EM maps in meshes, supporting that the model was built correctly.

**Extended Data Figure 4.**
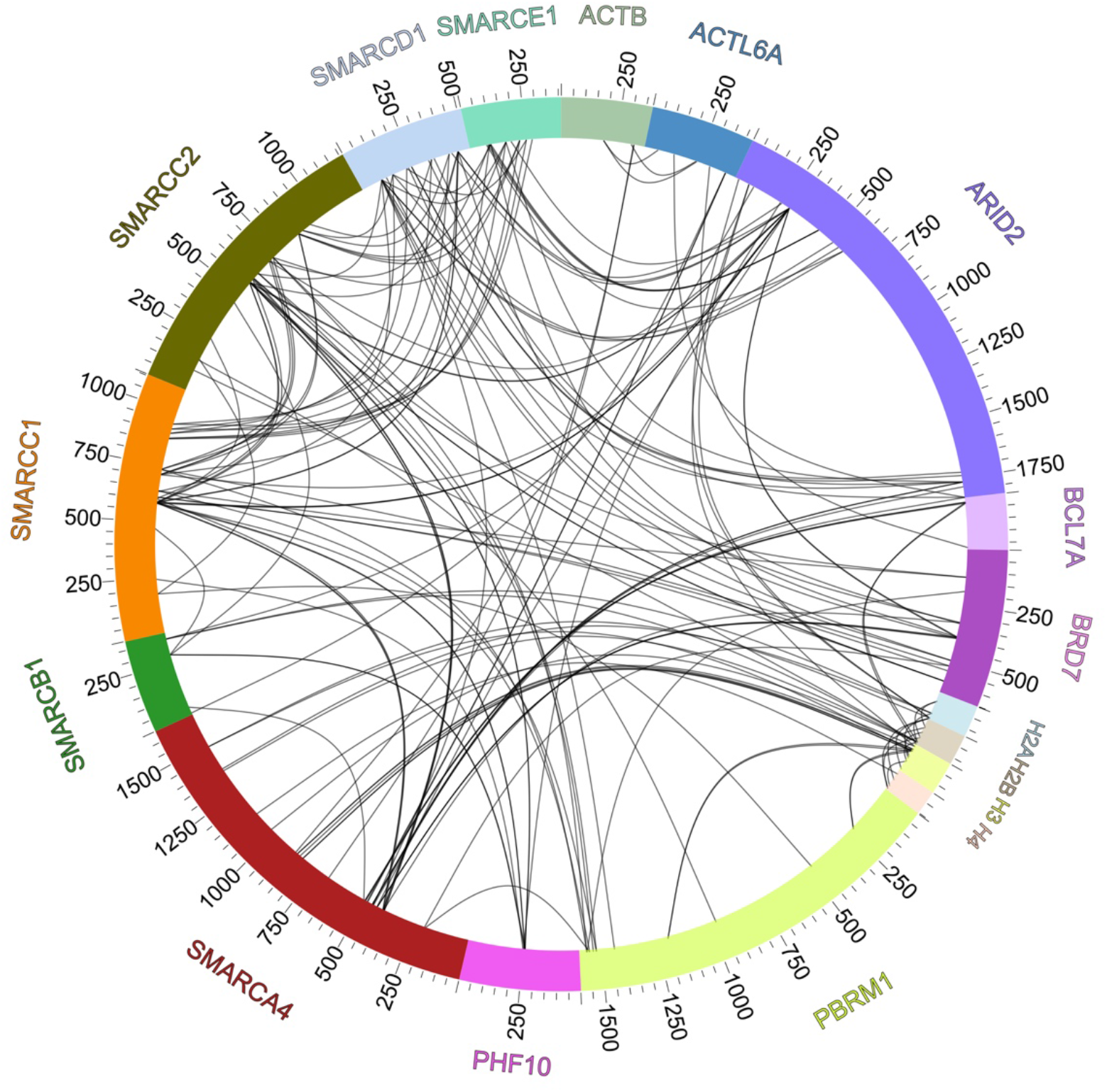
Cross-linking mass spectrometry. Schematic representation of inter-subunit cross-links within the PBAF-NCP complex in the presence of ADP-BeF_3_. Intramolecular cross-links were omitted for simplicity.

**Extended Data Figure 5.**
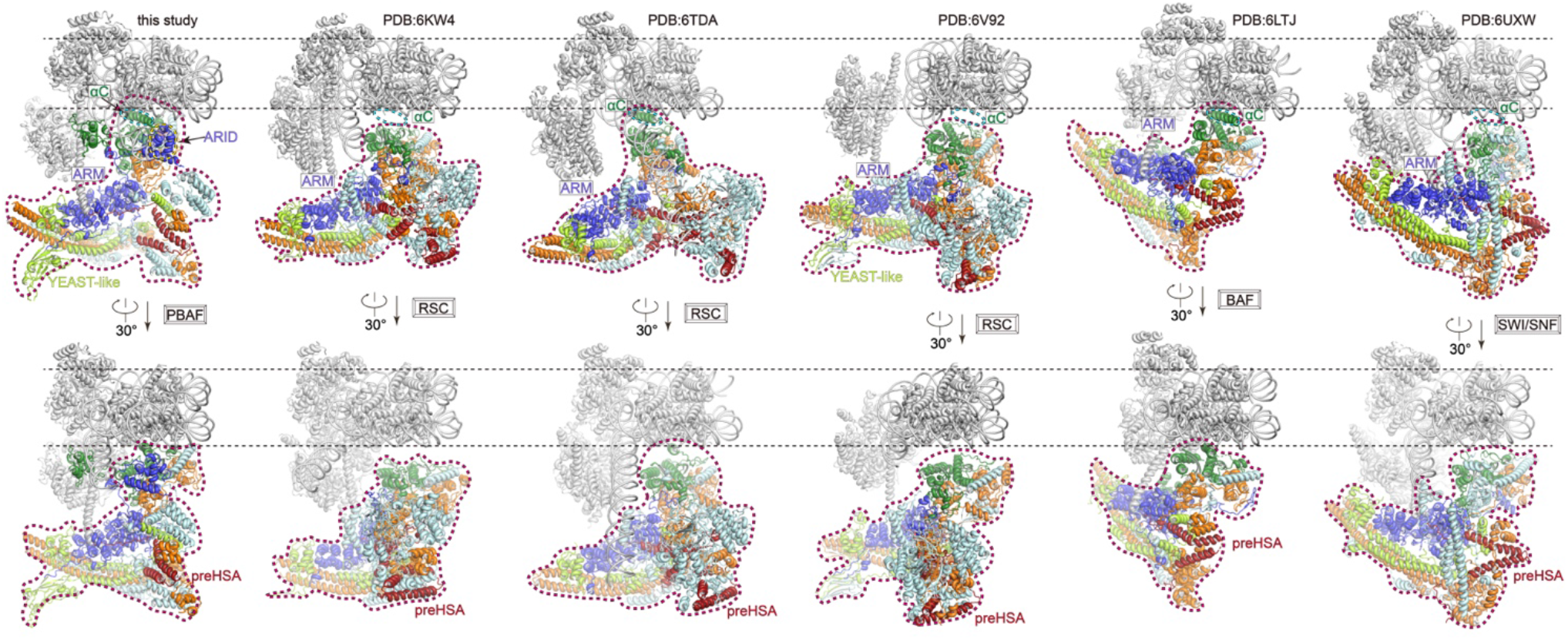
Structural comparison of nucleosome-bound human PBAF and other SWI/SNF family complexes. Two different views of the structures of nucleosome-bound human PBAF (this study), yeast RSC (PDB ID: 6KW4) (Ye et al., 2019), yeast RSC (PDB ID: 6TDA) (Wagner et al., 2020), yeast RSC (PDB ID: 6V92) (Patel et al., 2019), human BAF (PDB ID: 6LTJ) (He et al., 2020), and yeast SWI/SNF (PDB ID: 6UXW) (Han et al., 2020). The structures are shown with nucleosome in a similar orientation for comparison. Equivalent subunits are colored in the same color scheme.

**Extended Data Figure 6.**
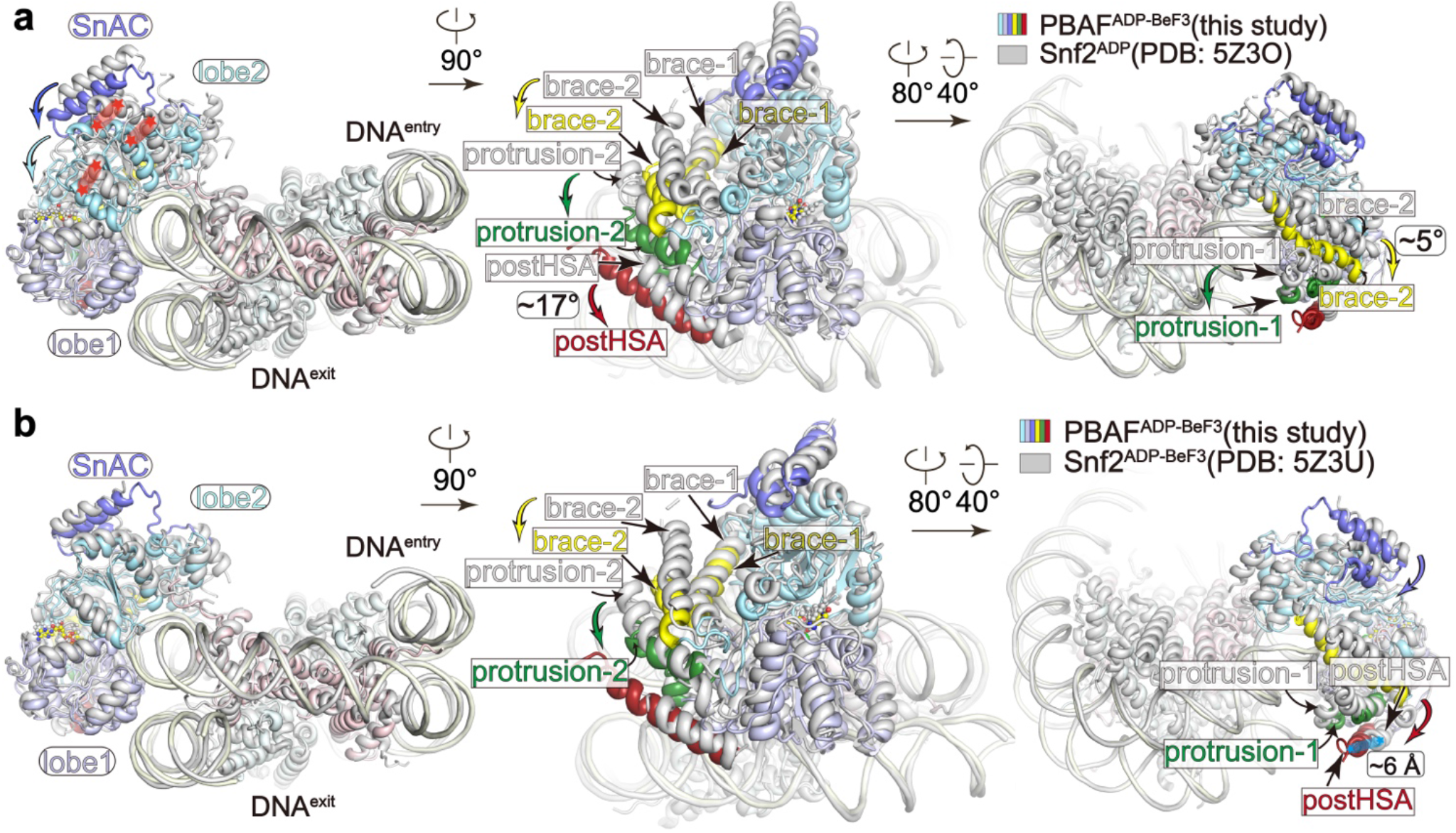
Nucleosome-bound ATPase in PBAF complex and isolated Snf2. Structural comparison of the nucleosome-bound ATPase in PBAF complex (ADP-BeF_3_-bound) and isolated Snf2 ATPase (Li et al., 2019) in the ADP-bound (**a**) and ADP-BeF_3_-bound (**b**) states, respectively. The structures are shown with nucleosome superimposed with structural differences indicated with arrows. Structure of Snf2-NCP is colored in grey and that of PBAF-NCP is colored as indicated.

**Extended Data Figure 7.**
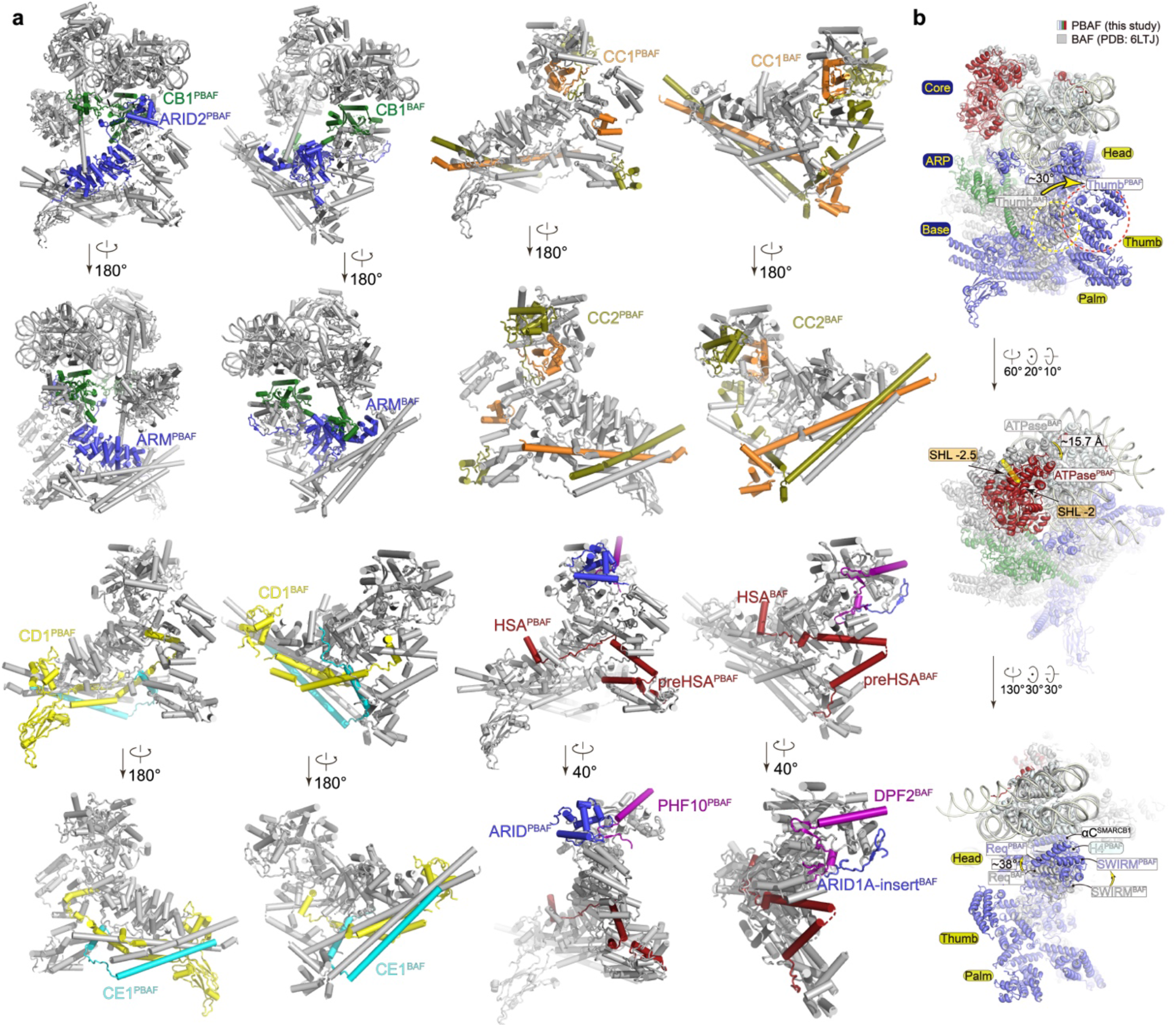
Structural differences in PBAF-NCP and BAF-NCP. (**a**) Structural comparison of PBAF-NCP (left panels) and BAF-NCP (right panels) structures with each subunit highlighted for comparison. (**b**) Superimposition of the two structures in three different views. Structural differences are indicated with arrows.

**Extended Data Figure 8.**
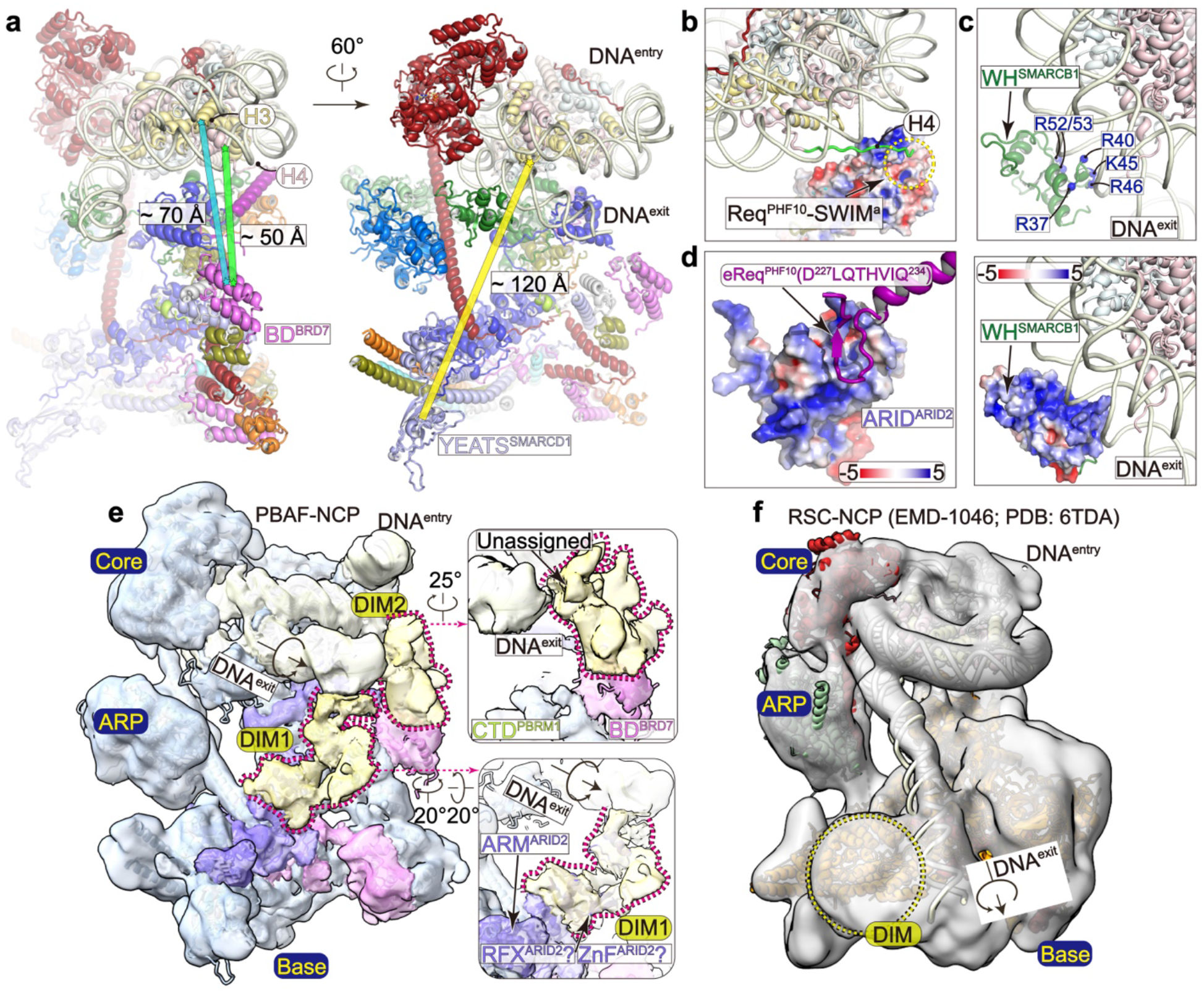
Positions of histone/DNA-binding domains in PBAF-NCP. (**a**) Overall structural model of PBAF-NCP in two different views. Distances between the bromodomain of BRD7 and histone fold domains of the nearest histone H3 and H4 are indicated in the left panel. Distance between YEATS-like domain of SMARCD1 and histone octamer is indicated in the right panel. (**b**) Close-up view of the contacts between histone H4 tail (shown in cartoon) and Req^PHF10^-SWIRM^SMARCC^ dimer. The acidic surface of Req^PHF10^-SWIRM^SMARCC^ is shown in electrostatic potential surface. (**c**) Close-up view of the interaction between WH^SMARCB1^ and nucleosomal DNA. Positively charged residues of WH^SMARCB1^ are indicated with blue balls in the upper panel. Positively charged surface of WH^SMARCB1^ is indicated with electrostatic potential surface in the lower panel. (**d**) Interaction between ARID^ARID2^ and PHF10. (**e**) Cryo-EM map of PBAF-NCP at low threshold shows unassigned regions of DIM1 and DIM2 that bind extranucleosomal DNA. Predicted structural model of RFX domain of ARID2 could be placed in the density of DIM1. (**f**) Cryo-EM map of RSC-NCP shows interaction between an unassigned DIM region and extranucleosomal DNA.

**Extended Data Figure 9.**
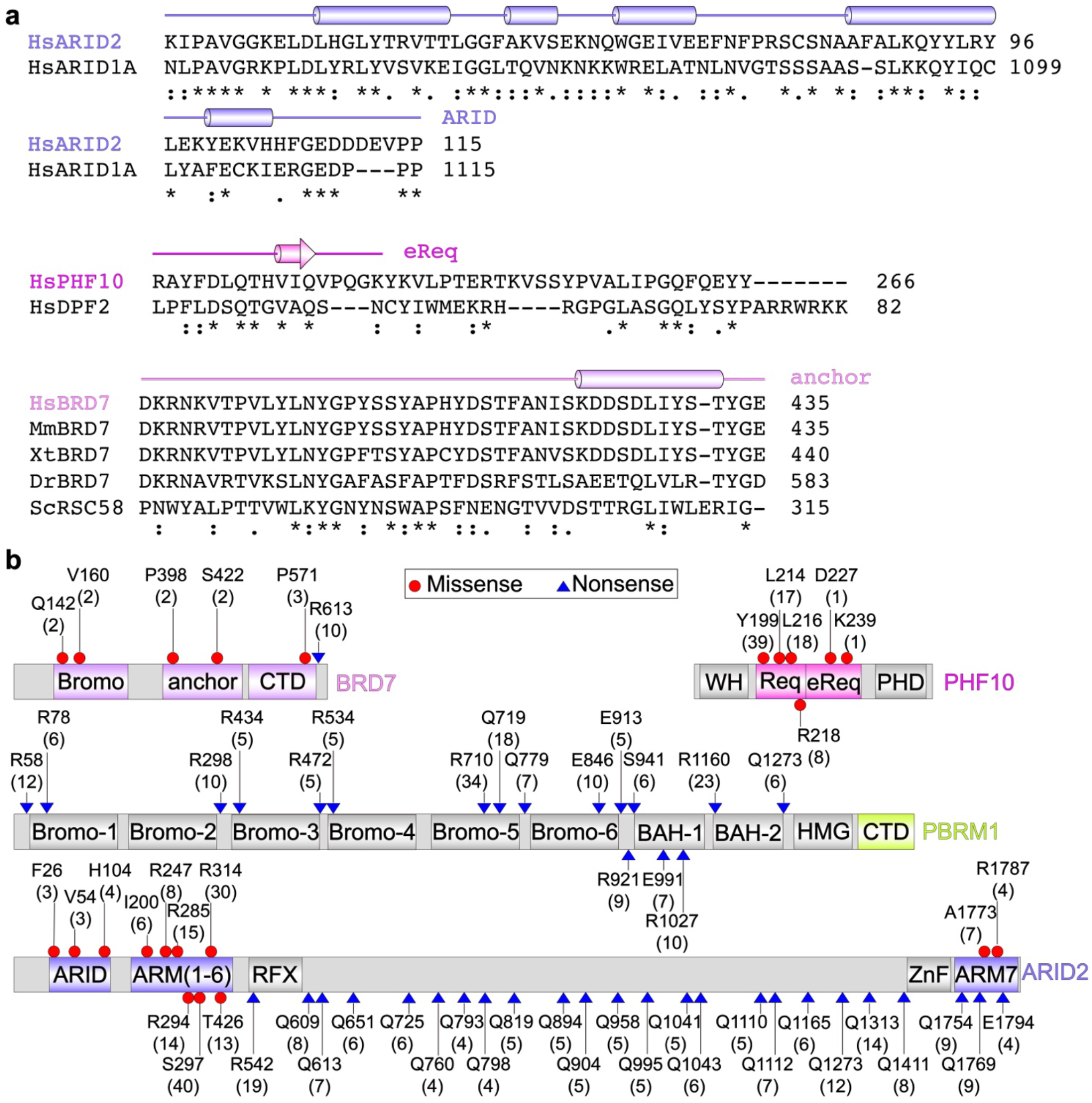
Sequence analysis of PBAF-specific organization and cancer-derived mutations. (**a**) Sequence analyses of PBAF-specific subunits with critical regions shown for comparison. (**b**) Cancer-derived mutations of PBAF-specific subunits.

**Extended Data Table 1.** Statistics of cryo-EM data collection and refinement.

|  | PBAF-NCP<br>(EMDB-XX)<br>(PDB-XX) | Base Module | ARP Module | Core Module |
| --- | --- | --- | --- | --- |
| **Data collection and processing** |  |  |  |  |
| Magnification | 64,000 x | 64,000 x | 64,000 x | 64,000 x |
| Voltage (kV) | 300 | 300 | 300 | 300 |
| Electron exposure ($e^-/\text{\AA}^2$) | 50 | 50 | 50 | 50 |
| Defocus range ($\mu\text{m}$) | -1.0 ~ -2.5 | -1.0 ~ -2.5 | -1.0 ~ -2.5 | -1.0 ~ -2.5 |
| Pixel size ($\text{\AA}$) | 1.334 | 1.334 | 1.334 | 1.334 |
| Symmetry imposed | C1 | C1 | C1 | C1 |
| Initial particle images (no.) | 2,071,055 | 2,071,055 | 2,071,055 | 2,071,055 |
| Final particle images (no.) | 43,759 | 31,995 | 31,995 | 43,759 |
| Map resolution | 4.4 | 4.1 | 5.4 | 3.4 |
| FSC threshold | 0.143 | 0.143 | 0.143 | 0.143 |
| Map resolution range ($\text{\AA}$) | 4.0-8.0 | 4.0-8.0 | 5.0-9.0 | 3.0-7.0 |
| **Refinement** |  |  |  |  |
| Model resolution ($\text{\AA}$) | 4.43 | 4.06 | 5.36 | 3.37 |
| FSC threshold | 0.143 | 0.143 | 0.143 | 0.143 |
| Model composition |  |  |  |  |
| Non-hydrogen atoms | 43,226 | 17,473 | 6,462 | 18,038 |
| Protein residues | 4,581 | 2,205 | 815 | 1,402 |
| Nucleotide residues | 330 | - | - | 330 |
| Ligands | 1× ADP,<br>1× Mg,<br>1× BeF<sub>3</sub> | - | - | 1× ADP,<br>1× Mg,<br>1× BeF<sub>3</sub> |
| B factors ($\text{\AA}^2$) | | | | |
| Protein | 104.09 | 183.4 | 107.69 | 57.50 |
| Nucleotide | 90.46 | - | - | 90.46 |
| Ligand | 20.04 | - | - | 20.04 |
| R.m.s deviations |  |  |  |  |
| Bond lengths ($\text{\AA}$) | 0.004 | 0.004 | 0.003 | 0.003 |
| Bond angles ($^\circ$) | 0.850 | 0.876 | 0.901 | 0.552 |
| Validation |  |  |  |  |
| MolProbity score | 1.93 | 1.84 | 1.84 | 1.73 |
| Clash score | 11.22 | 7.85 | 10.12 | 10.65 |
| Poor rotamers (%) | 0.31 | 0.05 | 0.00 | 0.51 |
| Ramachandran plot |  |  |  |  |
| Favored (%) | 94.79 | 93.73 | 95.48 | 96.93 |
| Allowed (%) | 5.07 | 6.27 | 4.52 | 3.07 |
| Disallowed (%) | 0.13 | 0.00 | 0.00 | 0.00 |

**Supplementary Video 1**

Composite cryo-EM map and structural model of PBAF-NCP.

**Supplementary Video 2**

Cryo-EM map of PBAF-NCP at low threshold showing unassigned regions of DIM1 and DIM2.

**Supplementary Video 3**

Structural comparison of PBAF-NCP (colored) and BAF-NCP (grey) with nucleosome superimposed.

